# The small RNA landscape is stable with age and resistant to loss of dFOXO signaling in *Drosophila*

**DOI:** 10.1101/2022.08.12.503791

**Authors:** Siobhan Gartland, Michael T. Marr

## Abstract

Aging can be defined as the progressive loss of physiological homeostasis that leads to a decline in cellular and organismal function. In recent years, it has become clear that small RNA pathways play a role in aging and aging related phenotypes. Small RNA pathways regulate many important processes including development, cellular physiology, and innate immunity. The pathways illicit a form of posttranscriptional gene regulation that relies on small RNAs bound by the protein components of the RNA-induced silencing complexes (RISCs), which inhibit the expression of complementary RNAs. In *Drosophila melanogaster,* Argonaute 1 (Ago1) is the core RISC component in microRNA (miRNA) silencing, while Argonaute 2 (Ago2) is the core RISC component in small interfering RNA (siRNA) silencing. The expression of Ago1 and Ago2 is regulated by stress response transcription factor Forkhead box O (dFOXO) increasing siRNA silencing efficiency. dFOXO plays a role in multiple stress responses and regulates pathways important for longevity. Here we use a next-generation sequencing approach to determine the effects of aging on small RNA abundance and RISC loading in male and female *Drosophila.* In addition, we examine the impact of the loss of dFOXO on these processes. We find that the relative abundance of the majority of small RNAs does not change with age. Additionally, under normal growth conditions, the loss of dFOXO has little effect on the small RNA landscape. However, we observed that age affects loading into RISC for a small number of miRNAs.

## Introduction

There are two major small RNA pathways in the somatic tissue of *Drosophila,* each of which have distinct functions and small RNA content [1, 2]. These pathways can be differentiated in *Drosophila* by the argonaute component present in the RNA-induced silencing complex (RISC). One of these is the microRNA (miRNA) pathway which is essential for cellular physiology and development [3]. The core protein component of the miRNA RISC is Argonaute1 (Ago1), which binds directly to miRNAs expressed from the genome. The other pathway is the small interfering RNA (siRNA) pathway, containing its core RISC protein Argonaute2 (Ago2), and acting as an important line of defense against RNA viruses in *D. melanogaster* [4–6]. In this pathway, double-stranded RNA from endogenous or exogenous sources is cleaved by Dicer-2 into siRNA duplexes loaded into Ago2 RISC.

The forkhead box O (FOXO) family of transcription factors responds to multiple cellular stresses, including nutrient deprivation, oxidative damage, heat shock, bacterial infection, and viral infection [7–9]. FOXOs are conserved from *Caenorhabditis elegans* to humans [7]. Whereas mammals possess four paralogs of FOXO, invertebrates have a single FOXO gene, making them valuable models for understanding the role of FOXO in gene regulation [7, 10, 11]. When the FOXO pathway is active in *C. elegans* and *D. melanogaster* lifespan is extended [11–13] and when FOXO is disrupted in *Drosophila* lifespan is reduced [14]. In *D. melanogaster,* dFOXO upregulates genes essential for the RNA interference pathway, including Ago1, Ago2, and Dicer-2 and has been shown to increase the efficiency of siRNA silencing [9]. The siRNA pathway also targets transposon expression in somatic tissue. Several studies in *Drosophila* have shown that the mRNA expression of many transposons increase with age and there is an ongoing debate over whether this increased expression is a driver or a byproduct of aging [15–18]. In any case, increased expression of Dicer-2 has been shown to increase lifespan and inhibit age-associated increases in transposon mRNA [16]. In addition, artificially inducing the *gypsy* transposon has been shown to shorten lifespan [19]. Therefore, the effect of dFOXO on the abundance of transposon siRNAs in RISC across age is of particular interest. The known targets of dFOXO in the miRNA pathway include only one RISC component, Ago1, while in the siRNA pathway dFOXO targets include both an upstream processing enzyme and a RISC component, Dicer-2 and Ago2 [9]. Thus, small RNA loading into each specific RISC may change over the fly lifespan and may be significantly impacted by dFOXO.

Because of the extensive interplay between small RNA pathways, dFOXO, and aging, we investigated the changes in small RNA abundance with respect to age in wildtype or dFOXO-null flies. Previous work investigating small RNAs and aging via RISC bound RNA sequencing showed that miRNAs’ loading into Ago2 increases with age [20]. However, these experiments were limited to males and did not interrogate the total small RNA repertoire for comparison. Here we investigate how total small RNAs, as well as RISC-bound small RNAs, change with age in male and female flies. In addition, we performed the same experiments in dFOXO-null animals to determine the role of the FOXO pathway on small RNA abundance and loading. In agreement with the work in male flies, we find that transposon siRNA abundance in Ago2 RISC decreases with age, whereas the levels of miRNA loading in Ago2 RISC increase with age. Loss of the FOXO pathway showed no detectable influence on this effect. We also find that deletion of dFOXO changes the abundance of four abundant miRNAs in females. In addition, we find that many miRNAs that vary in abundance in the whole animal with age do not exhibit a similar change in in abundance within miRNA RISC.

## Results

### The abundance of most miRNAs does not change with age in total small RNA

We sequenced two biological replicates of total small RNAs in whole wildtype (w^DAH^) adult flies at two timepoints, young (5-6 days) and old (30-31 days), to determine how the small RNA landscape changes with age. To separate miRNAs and siRNAs from piRNAs, we selected our reads to 19 to 25 bases in-silico. We mapped the libraries to a database containing the known miRNAs and transposons present in *Drosophila* and averaged the counts in our two replicates. We normalized the libraries by reporting each of our small RNA populations as a percent of total mapped reads. The 20 most abundant miRNAs in our young wildtype total small RNA libraries represent 79 to 94 percent of the reads that mapped to miRNAs (S1 and S2 Tables). With a stringent two-fold change set as a threshold, we find that 2 of the top 20 miRNAs change by increasing with age in wildtype males and females: miR-34-5p and miR-14- 3p (Fig 1). The observed increase in abundance of miR-34-5p with age is consistent with previous work [20, 21]. While no miRNAs decreased in abundance in male flies with age, two miRNAs decreased in female flies: miR-318-p and miR-994-5p (Fig 1A). miR-318 and miR-994 are positioned in close proximity in the genome, are abundantly expressed in ovaries, and are likely coregulated [22–24].

**Fig 1.**
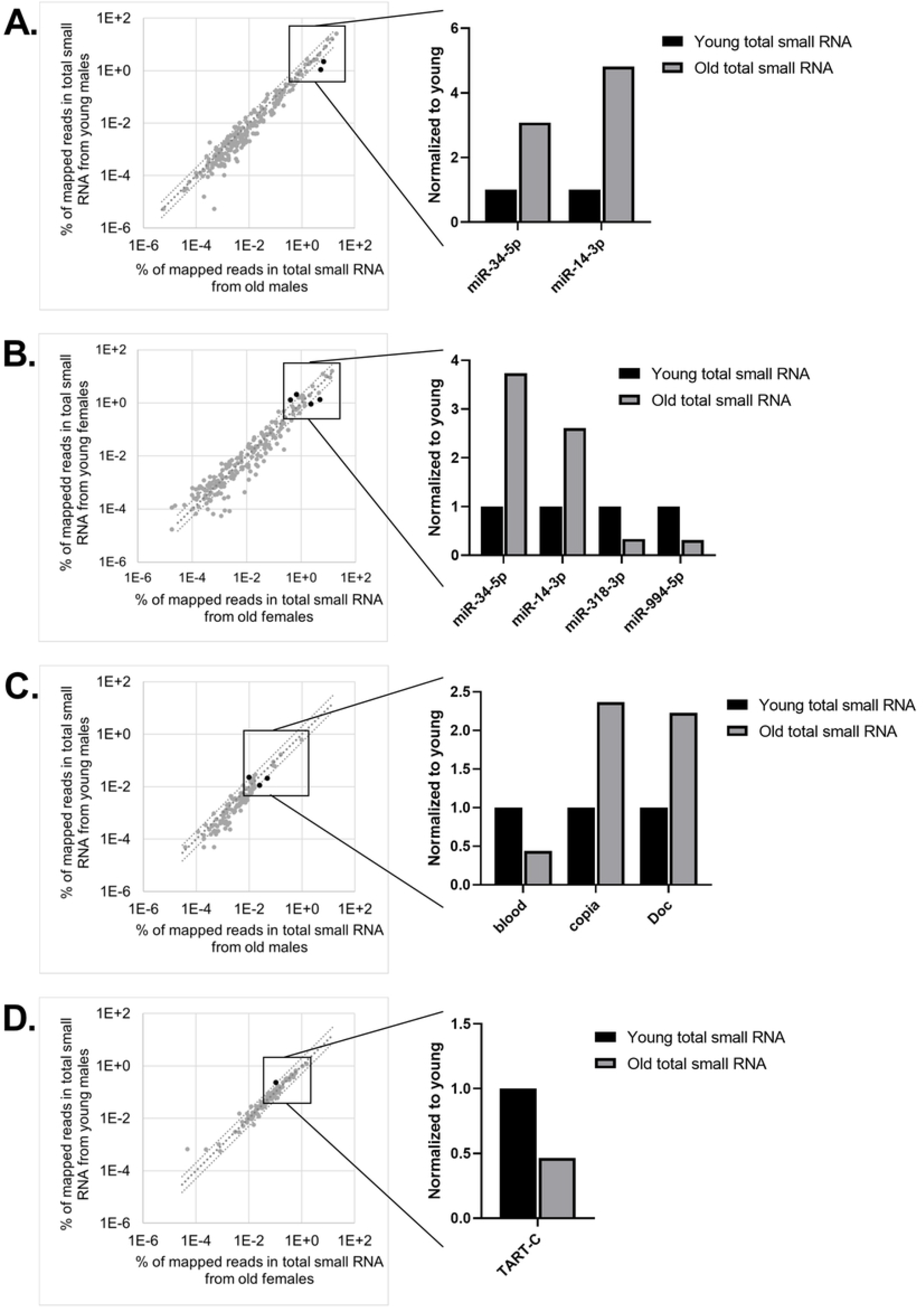
The abundance of most miRNAs and transposon siRNAs is stable with age. (A) Small RNAs from whole wildtype fly lysate were sequenced, mapped to known *Drosophila* miRNAs and transposons, and normalized by total mapped reads (n = 2). miRNA abundance is plotted as percent of reads mapped miRNAs and transposons in young (5-6 day old) and old (31 day old) males and (B) females (n = 2). Plotted in log10 scale. Top 20 miRNAs are boxed and top miRNAs that changed more than 2-fold are shown in orange. (C) siRNA abundance from males and (D) females.

### The abundance of most transposon siRNAs does not change across age in total small RNA

In order to specifically examine the transposon siRNAs produced by Dcr2, we conducted our siRNA analysis on 21 base reads [2, 25]. We focused on the 20 transposons with the most siRNAs mapped against them in young animals, representing 54 to 88 percent of reads mapped to transposons in our total small libraries. We found that in wildtype males, the relative abundance of siRNAs mapped against two transposons significantly increased by 2-fold or greater with age: *copia* and *Doc*. Conversely, the relative abundance of siRNAs mapped against the *blood* transposon decreased with age (Fig 1C). In females, *TART-C* was the only transposon that showed a significant age-dependent change in siRNA relative abundance with age, decreasing just over two-fold (Fig 1D).

### Preparation of polyclonal antibodies for Ago1 and Ago2 immunoprecipitation

In order to immunoprecipitate the RISC complexes, we generated polyclonal antiserum to Ago1 and Ago2 (Fig 2A). Glutathione-S-transferase (GST) fusions to the N-terminus of Ago1 (aa 1-300) or Ago2 (aa 1-490) were purified from *E. coli* and used to immunize guinea pigs. The antisera were screened for their ability to immunoprecipitate either Ago1 or Ago2. Preimmune or immune sera was used to precipitate Ago1 or Ago2 from whole fly lysates. The precipitated proteins were probed with either a second guinea pig antisera or a commercial antibody against Ago1 or Ago2 (Fig 2B and C). Comparison with a commercial anti-Ago1 antibody indicates that the guinea pig antisera effectively and specifically immunoprecipitates Ago1; likewise, the anti- Ago2 antisera effectively and specifically immunoprecipitates Ago2 and is able to deplete Ago2 from lysate (S1 Fig).

**Fig 2.**
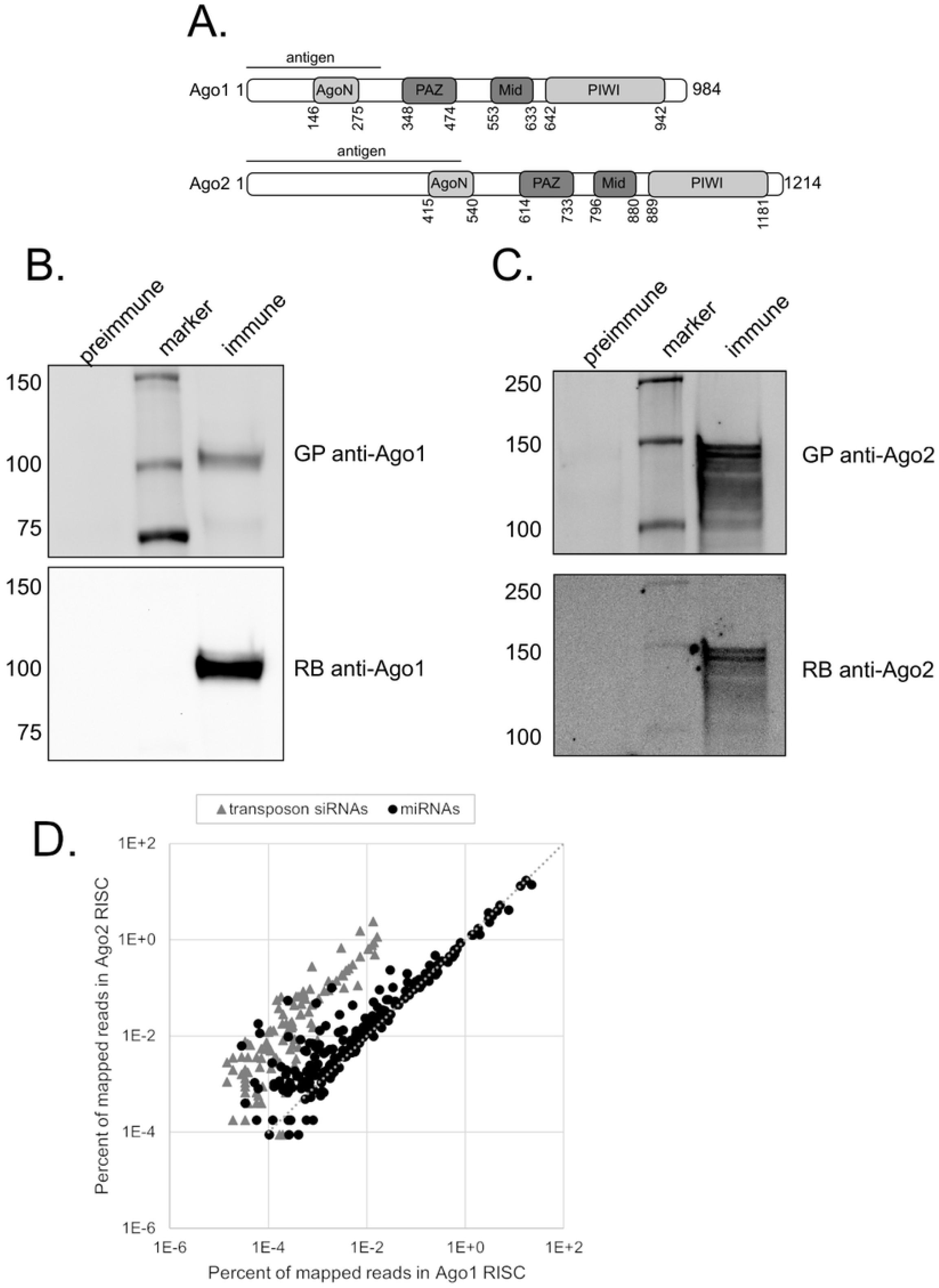
Polyclonal antibodies immunoprecipitate Ago1 and Ago2 exclusively. (A) Domain map of Ago1 and Ago2. (B) The guinea pig (GP) antiserum against Ago1 is able to immunoprecipitate Ago1 as detected by a commercial rabbit (RB) anti-Ago1 antibody [Abcam]. (C) The guinea pig (GP) antiserum against Ago2 is able to immunoprecipitate Ago2 as detected by a commercial rabbit (RB) anti-Ago2 antibody [Abcam]. (D) Small RNAs immunoprecipitated with Ago1 or Ago2 antisera from whole wildtype male lysate were sequenced and mapped to known *Drosophila* miRNAs and transposons (n = 2). miRNAs (black circles) and siRNAs (gray triangles) are shown as percent of total mapped reads. Several miRNAs that have been previously observed to be highly abundant in Ago2 RISC are noted. Representative plot.

### The abundance of most miRNAs in Ago1 RISC is stable with age

To investigate whether aging affects small RNAs’ loading into RISC, we immunoprecipitated the argonaute proteins from whole fly lysates and made small RNA sequencing libraries. This allowed us to construct libraries specifically from the small RNAs bound to endogenous Ago1 and Ago2. Then, two biological replicates of each were generated and these libraries were mapped to miRNAs and transposons, in the same manner as the total small RNA libraries. Overall, the results are consistent with previous observations [26], siRNAs make up a higher percentage of small RNAs in the Ago2 RISC and miRNAs are preferentially loaded in the Ago1 RISC (Fig 2D). Additionally, we also observed differential loading of a few miRNAs, which are more abundant in Ago2 RISC, such as miR-988-5p, miR-2a-2p, and let-7- 3p, (Fig 2D), consistent with previous findings[2].

For the Ago1 RISC, we focused our analysis on the same 20 most abundant miRNAs as we did in the total small RNA libraries. As expected from the total small RNA library result, the abundance of most of the top 20 miRNAs found in RISC remains stable with age (Fig 3A and 3B). However, the two miRNAs which show an increase in abundance in total small RNA had different outcomes in terms of RISC loading. In males, Ago1 RISC miR-34-5p abundance increases with age proportional to its increase in total abundance (Fig 3A). However, in females, miR-34-5p loading does not reflect its increase in the total small RNA pool (Fig 3B). Additionally, the increased abundance with age seen for miR-14-3p in the total small libraries is not reflected in the Ago1 RISC libraries for either sex (Fig 3A and B).

**Fig 3.**
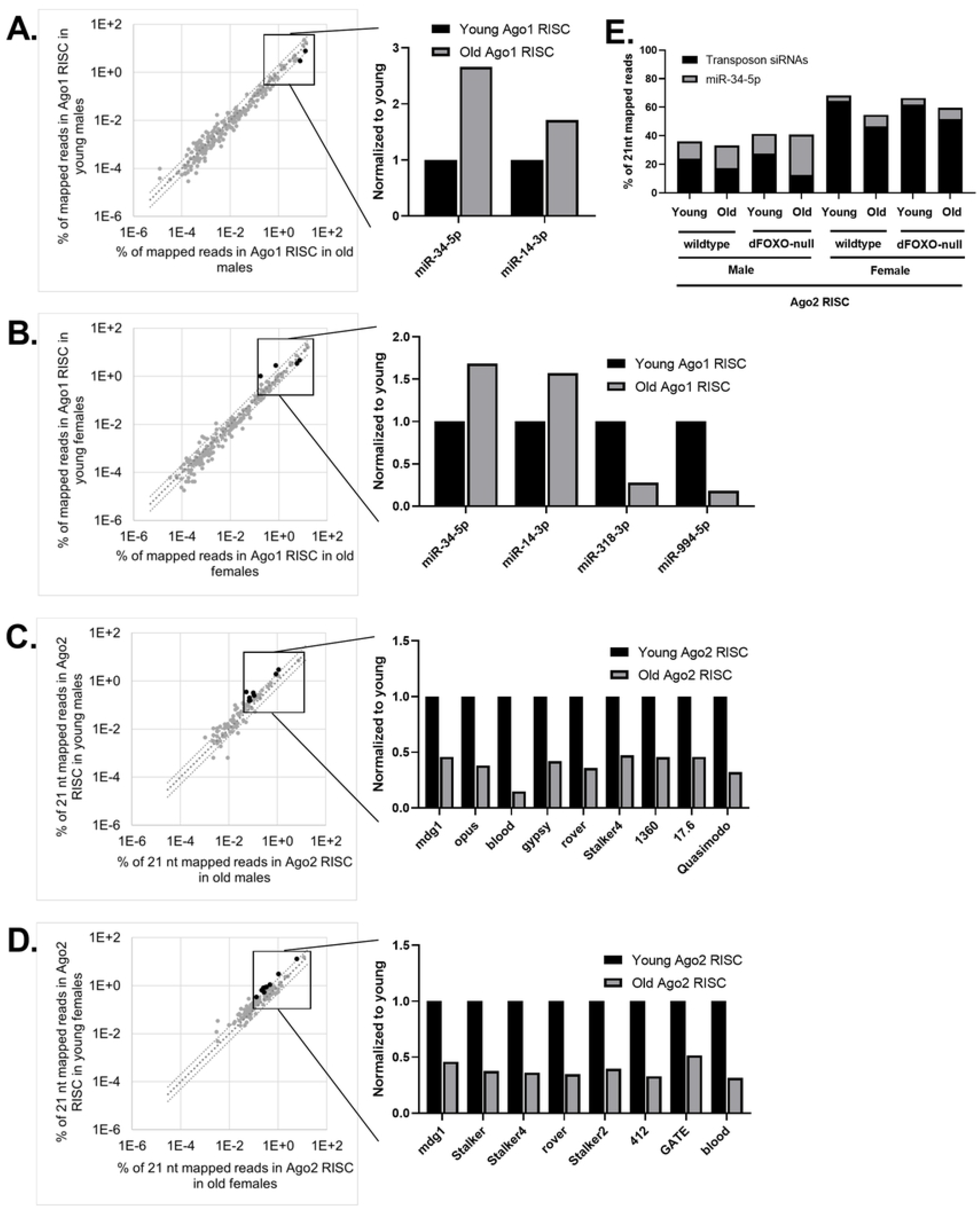
Most miRNAs in Ago1 RISC are stable with age whereas transposon siRNAs in Ago2 RISC decrease with age. Small RNAs immunoprecipitated with Ago1 RISC or Ago2 RISC from young (5 day old) or old (35 day old) whole flies were sequenced and mapped to known *Drosophila* miRNAs and transposons. (A) miRNAs in Ago1 RISC from old and young males are plotted as percentage of mapped reads. miR-34-5p increases with age by more than 2-fold while miR-14-3p does not. (B) miRNAs in Ago1 RISC from old and young females are plotted as percentage of mapped reads. miR-34-5p and miR-14-3p do not change by 2-fold. miR-318-3p and miR-994-5p decrease by 2-fold with age. (C) Small RNA libraries were size selected to 21 nt in silico and mapped to miRNAs and transposons in males and (D) females. In both sexes several of the most abundant transposons decrease abundance in Ago2 RISC with age.

### The abundance of transposon siRNA decreases with age in Ago2 RISC

To investigate the changes in Ago2 RISC, we immunoprecipitated Ago2 and made small RNA libraries from the precipitated material. As expected from previous work [1], siRNAs mapped to transposons make up a larger percentage of the Ago2 libraries than the Ago1 libraries (S2 Fig). In the Ago2 RISC libraries, the abundance of siRNAs mapped to the top transposons in males and females either decreased with age or did not change (Fig 3B). Of the transposon siRNAs that showed a decreased overall abundance with age, the Ago2 loading of siRNAs of nine transposons decreased in old male and eight decreased in old female flies (Fig 3C and 3D). Taken as a whole, transposon siRNA abundance decreases with age in Ago2 RISC, in both males and females. Additionally, this difference is associated with an increase of the 21 nt long form of miR-34-5p, as previously observed in males [20] (Fig 3E).

### dFOXO dependent changes in total small RNAs

We sought to determine if dFOXO signaling has an effect on the small RNA landscape. To this end, we made total small RNA Illumina libraries from flies with a large deletion in the dFOXO locus, removing a large portion of the coding region and creating a null allele (w^DAH^ Δ94) [14]. Otherwise, these animals are isogenic with w^DAH^. Interestingly, the effect of the loss of dFOXO is very complex. In general, under these growth conditions, the majority of the population of total small RNAs are unaffected by the disruption of the FOXO pathway. However, there are reproducible differences between wildtype and dFOXO-null animals, some of which are age or sex specific.

We first examined the effects of the loss of dFOXO on the two miRNAs that have the greatest age dependent change in wildtype animals, miR-14-3p and miR-34-5p. dFOXO-null males and females show the same pattern of abundance for total miR-14-3p. In young animals, miR-14-3p is lower relative abundance in dFOXO-null animals than in wild-type. However, it appears that miR-14-3p accumulates at a greater rate in dFOXO-null animals as the 35-day old flies in both genotypes report virtually the same amount of miR-14-3p (Fig 4A). By contrast, miR-34-5p shows a sex specific effect. In males, miR-34-5p abundance is equivalent in both genotypes (Fig 1A) but in females miR-34-5p is less abundant in dFOXO-null animals at both ages (Fig 4B).

**Fig 4.**
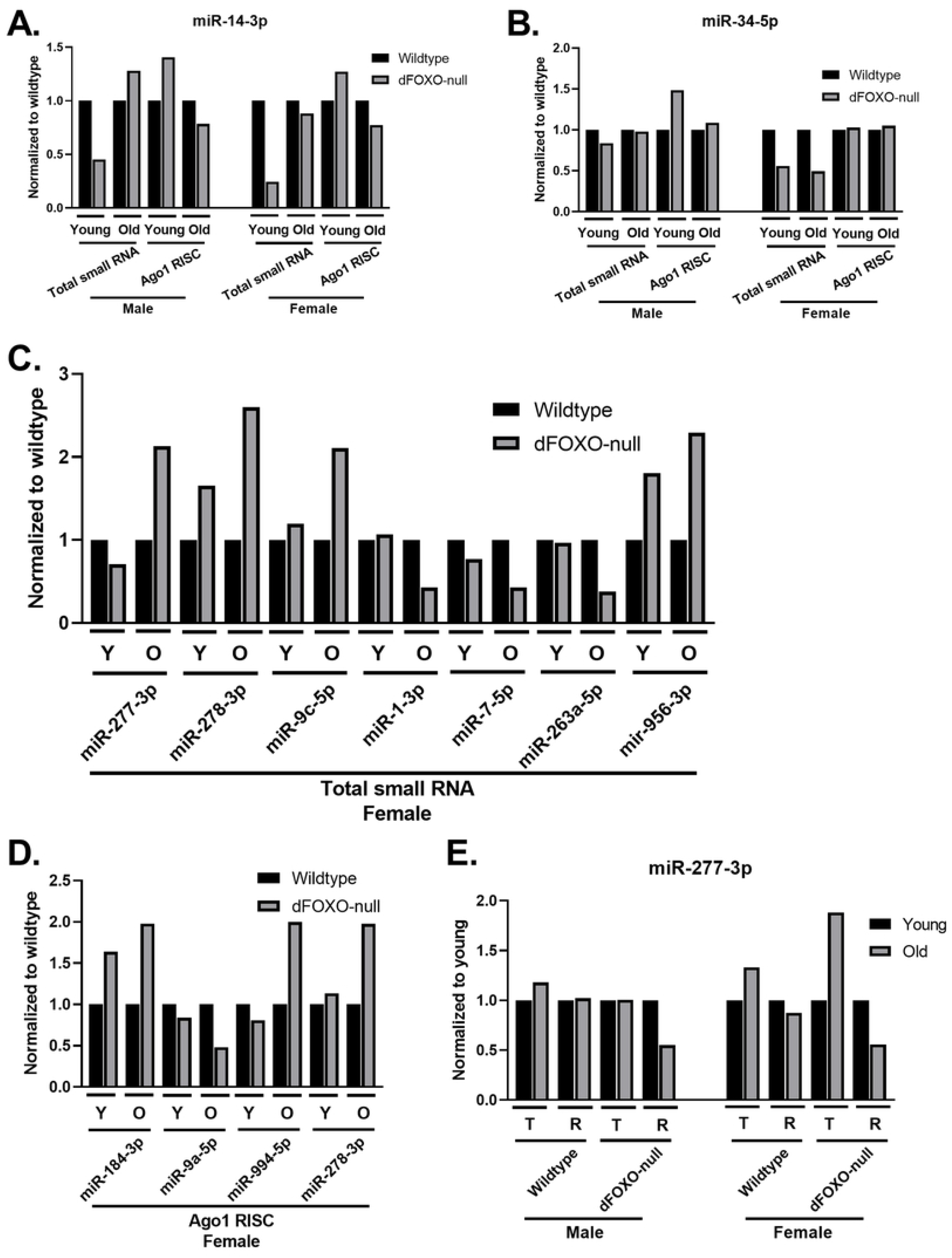
dFOXO dependent changes on small RNAs in total abundance and Ago1 RISC. Small RNAs from wildtype or dFOXO-null flies collected from whole lysate or Ago1 RISC were sequenced, mapped to known *Drosophila* miRNAs and transposons, and normalized by total mapped reads (n = 2). (A) miR-14-3p abundance from dFOXO-null flies in total small RNA or Ago1 RISC were normalized to wildtype. dFOXO-null animals have decreased miR-14-3p at young age in total abundance but not in Ago1 RISC. (B) miR-34-5p abundance in dFOXO-null females is half of wildtype in total abundance but similar abundance in Ago1 RISC. (C) In total small RNA in females, several miRNAs at young (Y) and old (O) age have differential abundance depending on dFOXO. (D) In Ago1 RISC in females, 3 different miRNAs than in total small RNA and miR-278-3p changed dependent on dFOXO. (E) miR-277-3p in total small RNA (T) and Ago1 RISC (R) has decreased abundance at old age only in Ago1 RISC of dFOXO-null males and females.

Other changes in abundance were observed in females only. Three miRNAs, miR-277- 3p, miR-278-3p, and miR-9c-5p, are more abundant in older dFOXO-null animals compared to older wildtype (Fig 4C). Three miRNAs are less abundant in older dFOXO-null animals than in older wildtype, miR-1-3p, miR-7p, and miR-263a-5p (Fig 4C). One miRNA, miR-956-3p, is more abundant in dFOXO-null animals than wildtype at both ages (Fig 4C).

### dFOXO dependent changes in Ago1 RISC

To investigate effects of the dFOXO deletion on miRNA loading, we immunoprecipitated Ago1 RISC in wildtype and dFOXO-null animals and compared miRNA abundance. In the young animals of both sexes, disruption of dFOXO had no significant effect, above our stringent 2-fold threshold, on Ago1 RISC loading despite the miRNA abundance differences seen in the total small RNA libraries. In older female dFOXO-null animals, four miRNAs showed increased in abundance in Ago1 RISC, while one miRNA showed a decrease in loading (Fig 4D). Of the four miRNAs that exhibited changes in Ago1 RISC loading, miR-184-3p and miR-9a-5p do not show a change in total abundance. One miRNA is affected by loss of dFOXO in both sexes in older animals; miR-277-3p is comparatively less abundant in Ago1 RISC in dFOXO-null animals (Fig 4E).

### The basal landscape of transposon siRNAs is dFOXO sensitive

As the identity of the most abundant transposon siRNAs differ between wildtype and the dFOXO-null animals, it is quite difficult to directly compare specific transposons. In both sexes of wildtype animals, *297* is the transposon with the most siRNAs. siRNA reads mapping to *297* are much less abundant in dFOXO-null libraries, with 4-fold fewer in females and 8-fold fewer in males (Fig 5A and B). The transposon with the most abundant siRNAs in dFOXO-null males is *roo* (S2 Table), which has 2-fold more siRNA abundance in wildtype males and but has a similar abundance in both female genotypes (Fig 5A and B).

**Fig 5.**
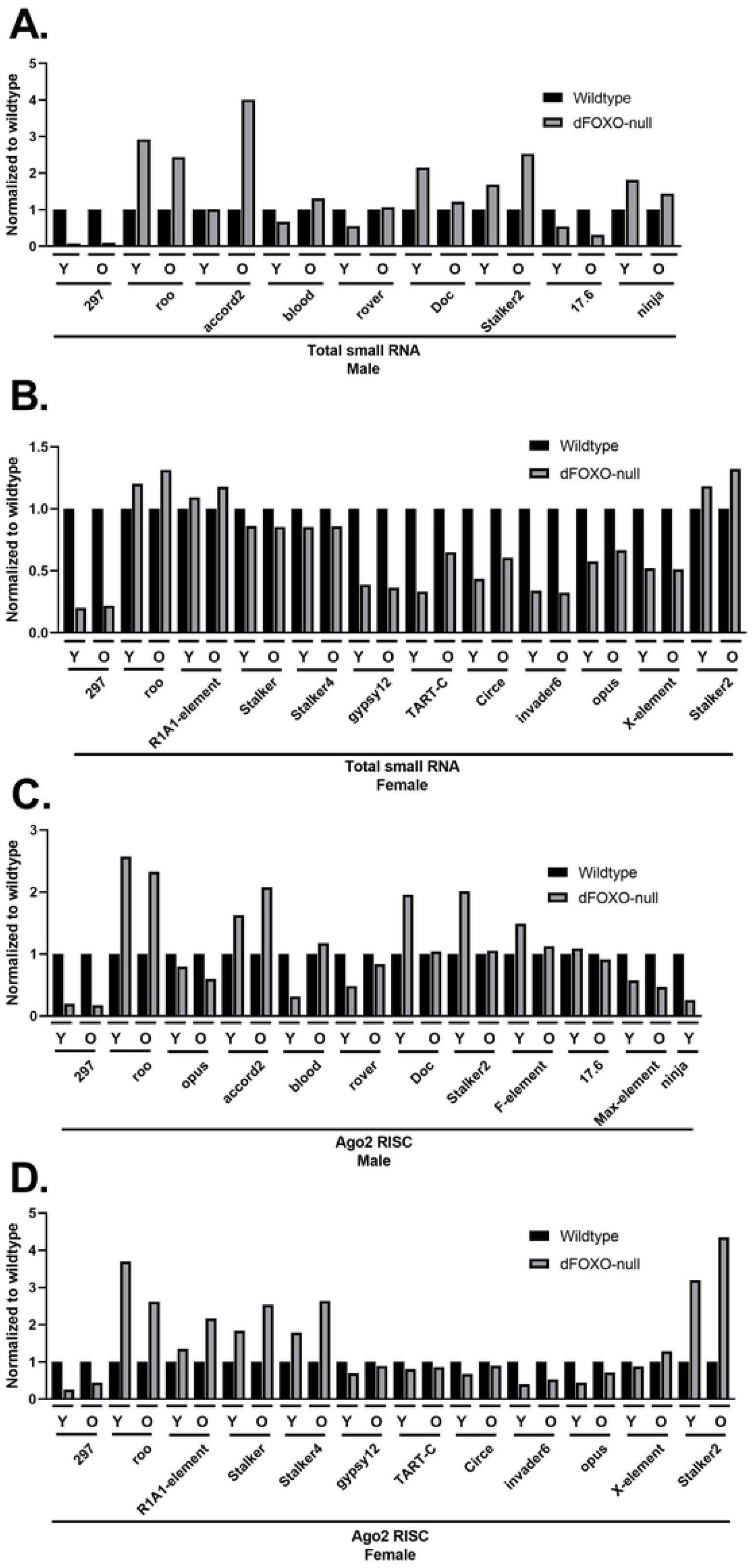
dFOXO dependent changes on transposon siRNAs in total abundance and Ago2 RISC. Small RNAs from wildtype or dFOXO-null flies collected from whole lysate and Ago2 RISC were sequenced, size selected to 21 nt in silico, mapped to known *Drosophila* miRNAs and transposons, and normalized by total mapped reads (n = 2). (A) Many transposon siRNAs in total small RNA at either or both young (Y) and old (O) age have differential abundance dependent on dFOXO in males and (B) females. (C) Most of these transposons also have differential abundance dependent on dFOXO in Ago2 RISC as well in males and (D) females. *Ninja* was not detected in the male wildtype Ago2 RISC old timepoint.

### dFOXO dependent changes in Ago2 RISC

The siRNA RISC occupancy of several transposons differs between wildtype and dFOXO-null animals. However, most of the differences can be traced to differences in total siRNA abundance. For example, in the cases of *297* and *roo*, transposons with the most abundant siRNAs in wildtype and dFOXO-null respectively, the difference in abundance is reflected in the siRNA content of Ago2 RISC (Fig 5C and D). In males there are two notable exceptions. First, in old animals, although the *stalker2* occupancy is the same for both genotypes, the total siRNA abundance is greater in dFOXO-nulls (Fig 5A and C). Second, while the total siRNAs matching *ninja* are similar between genotypes, *ninja* siRNAs are not loaded as efficiently into Ago2 RISC in wildtype animals as in the dFOXO-null animals (Fig 5A and C). There are also exceptions in females. For example, siRNAs matching *stalker, stalker2, stalker4, gypsy12, circe* and *TART-C* are overrepresented in the Ago2 RISC relative to their total abundance.

## Discussion

In wildtype *Drosophila* of both sexes, the relative abundance of most small RNAs did not change with age. This indicates that the small RNA landscape remains relatively stable in otherwise unstressed aging animals. However, there were a handful of miRNAs whose expression was affected by aging. Notably, the abundance of both miR-14-3p and miR-34-5p increases with age in male and female *Drosophila* (Fig 1A). Interestingly, these age dependent changes in the total small RNA population are not always reflected in Ago1 RISC, suggesting that loading into Ago1 does not always correlate with the total abundance of a given miRNA (Figs 1A, 1B, 3A, and 3B). This is true even for miRNAs, such as miR-14-3p, which are preferentially loaded into Ago1 rather than Ago2 [2]. The total abundance of miR-14-3p increases by more than 2-fold with age, but its abundance in Ago1 RISC does not increase at that same rate. This suggests that the rate of miR-14-3p loading into Ago1 RISC may decrease with age. Perhaps this corresponds to the metabolism dysregulation that older animals experience; as insulin production and normal fat metabolism are key functions of mir-14 [27, 28].

Previous studies have only investigated miRNA occupancy in RISC in males as a function of age [20], and no studies had investigated the total small RNA landscape across age for comparison. Here we find that aging females show an age dependent pattern in their total miRNA landscape that is similar to males, with major differences arising from ovary associated miRNAs, which decrease with age (Fig 1B). The age associated decreases of miR-318 and miR-994 were greater in wildtype than in dFOXO-null females (Fig 4C). This suggests that the ovaries of dFOXO-null animals are more similar to the ovaries of older wildtype animals and may play a role in the decreased fertility reported in the dFOXO-null genotype [14].

The loss of dFOXO, in normal growth conditions, leads to only small changes in the overall small RNA landscape with respect to age, with most of the effects limited to only one of the time points. miR-277-3p abundance increases in old animals to a much greater extent in dFOXO-null animals than in wildtype. Intriguingly, the constitutive expression of this miRNA has been shown to shorten lifespan and interfere with normal metabolism [29]. However, the loading of miR-277-3p into Ago1 RISC in both sexes of dFOXO-null animals is depressed at the old timepoint, even though miR-277-3p abundance is elevated in old dFOXO-null females compared to wildtype. This result suggests a complex dynamic between the total pool of small RNAs and the efficiency of their loading into RISC (Fig 4E). Therefore, both increased abundance and decreased loading of miR-277-3p could contribute to the shorter lifespan of dFOXO-null animals.

One exception to the general trend in age specific differences is miR-956-3p. This miRNA is more abundant in dFOXO-null females at both timepoints (Fig 4C). Interestingly, the knockout of miR-956-3p appears to be protective against the *Drosophila* C Virus [30]. Given dFOXO’s role in defending against RNA viruses [9], perhaps dFOXO does so by acting through a number of avenues, repressing the expression of miR-956-3p while increasing that of Ago2 and DCR-2. These differences in miRNA abundance and loading into RISC may have more to do with dFOXO as a transcription factor than its direct effects on the expression of miRNA machinery; we observed only changes to particular small RNAs, and not a global trend.

The siRNAs landscape targeting transposons differs between the 2 genotypes tested here. This may be due to the apparent difference in active transposons between these two strains [19]. However, the differences in siRNA abundance both in total small RNA population and loaded in RISC do not correlate with the differences seen in transposon mRNAs [19]. This suggests a complex interaction between transposon mRNA expression, siRNA formation and RISC loading that is still unexplored.

The most abundant transposon siRNAs loaded into Ago2 RISC trend downward with age (Fig 3C and D). As our analysis is based upon normalizing each library to the total miRNA and transposon siRNA reads mapped, this observation corresponds directly to an increase in the short isoforms of miRNAs in Ago2 RISC, which agrees with previous observations reported in males [20] (Fig 4E). This could reflect an age dependent decrease in Ago2 loading specificity that allows for the increased the silencing efficiency of miRNAs like miR-34-5p, which has an established role in preventing the aging of the brain [21]. However, the associated decrease in siRNAs occupancy in Ago2 RISC may leave the animal less able to defend against transposon activity as they age, potentially contributing to the age associated increase in transposon expression. Our results with total small RNAs and RISC bound small RNAs in both sexes largely complement what was previously found relating to aging and the Ago2 RISC complex. We concur that as the animal ages, there is an increase in miRNA loading into Ago2 RISC [20].

## Methods

### Fly Lines

w^Dah^ and *FOXO^Δ94^* were described [14].

### Antisera Generation

Glutathione-S-transferase (GST) fusions to the N-terminus of Ago1 (aa 1-300) or Ago2 (aa 1-490) were purified from *E. coli* and sent to Cocalico Biologicals, Inc. for guinea pig immunization and antisera exsanguination.

### Argonaute Immunoprecipitation

Forty whole flies were homogenized in 500 μL of chilled lysis buffer [31]. Lysates were incubated on ice for 10 minutes and centrifuged at 15,000 rpm for 15 minutes at 4 °C. Two hundred microliters of supernatant were kept as input. For the Ago1 immunoprecipitation, 4 μL of polyclonal guinea pig antibodies BRD-GP-3 and 4 μL BRD-GP-4 were added to 100 μL of lysate. For the Ago2 immunoprecipitation, the same was done with polyclonal guinea pig antibodies BRD-GP-5 and BRD-GP-6. Lysates were incubated with antibody and were rotated for 1 hour at 4 °C. Fifty microliters of 25% slurry Pierce protein A/G magnetic agarose [Thermo Fisher] were prepared for each IP. The magnetic agarose was washed in 1 mL of lysis buffer three times before being resuspended in the original volume. Forty microliters of slurry were aliquoted into new tubes, supernatant was removed, and the lysates with antibody were added. Lysates and magnetic agarose were incubated overnight with rotation at 4 °C. Agarose was washed five times for 5 minutes in 500 μL of lysis buffer at 4 °C before elution for RNA extraction or western blotting.

### RNA Extraction, RNA Size Selection, and Small RNA Sequencing

RNA was extracted from 40 whole flies or immunoprecipitated Ago1 or Ago2 using Trizol reagent [Invitrogen] following the manufacturer’s protocol. RNA extracted from whole flies was sized selected to 19 to 35 nt fragments using 17% 19:1 acrylamide:bisacrylamide urea gels [National Diagnostics]. Gels were sliced from 19-35 nt, crushed, and eluted overnight in 500 μL 0.3 M sodium acetate with rotation. Twenty micrograms of glycogen and small RNA spike ins [32] were added to the supernatant before it was precipitated with 750 μL isopropanol at −20 °C for 1 hour, centrifuged at 15,000 rpm for 30 minutes at 4 °C, washed in 80% EtOH, air dried, and resuspended in 50 μL H_2_O. Small RNA libraries were constructed using the NEBNext^®^ Small RNA Library Prep Set for Illumina^®^ [New England Biolabs] following the manufacturer’s protocol. Libraries were sequenced on a NextSeq500 platform [Illumina].

### Computational Analysis

Processing of sequence files was conducted using the Galaxy web platform [33] and the Mississippi server sponsored by the ARTbio bioinformatics facility of the Institut de Biologie Paris Seine based at the University Pierre & Marie Curie. Adapters were removed using the application Trim Galore! designed by Felix Krueger on the default settings and by inputting the adapter sequence GATCGTCGGACTGTAGAACTCTGAACGTGTAGATCTCGGTGGTCGCCGTATCATT. The trimmed sequence files were processed through the Reverse-Complement application. These trimmed and reverse complemented files were then size selected using Filter FASTQ [34] to filter out reads longer than 25 nt or filtered to only include 21 nt reads for transposon siRNA analysis. Reads were mapped to the dm6 genome or an in-house FASTA consisting of spike-ins [32], known *Drosophila* transposons, mature miRNAs, and miR* strands using sR_bowtie [35] set to match and report all valid alignments with 1 mismatch allowed. Alignments were quantified using featureCounts [36]. miR* strands, and transposons with an average of less than 5 reads across the libraries were removed from analysis to avoid noise using DEBrowser v1.18.2 [37]. Libraries were normalized by total mapped reads, adapted from Czech *et. al* 2008 [2, 26].

## Acknowledgements

We thank Meghan T. Harris, Zachary Knotts, William Dahl, Guilherme Gatti da Silva, and Kevin Clark for critical reading of the manuscript, Ildar Gainetdinov and the Zamore lab for technical advice and small RNA spike-ins, and Joyce Rigal for advice and support.

## Supporting Information

**S1 Fig.**
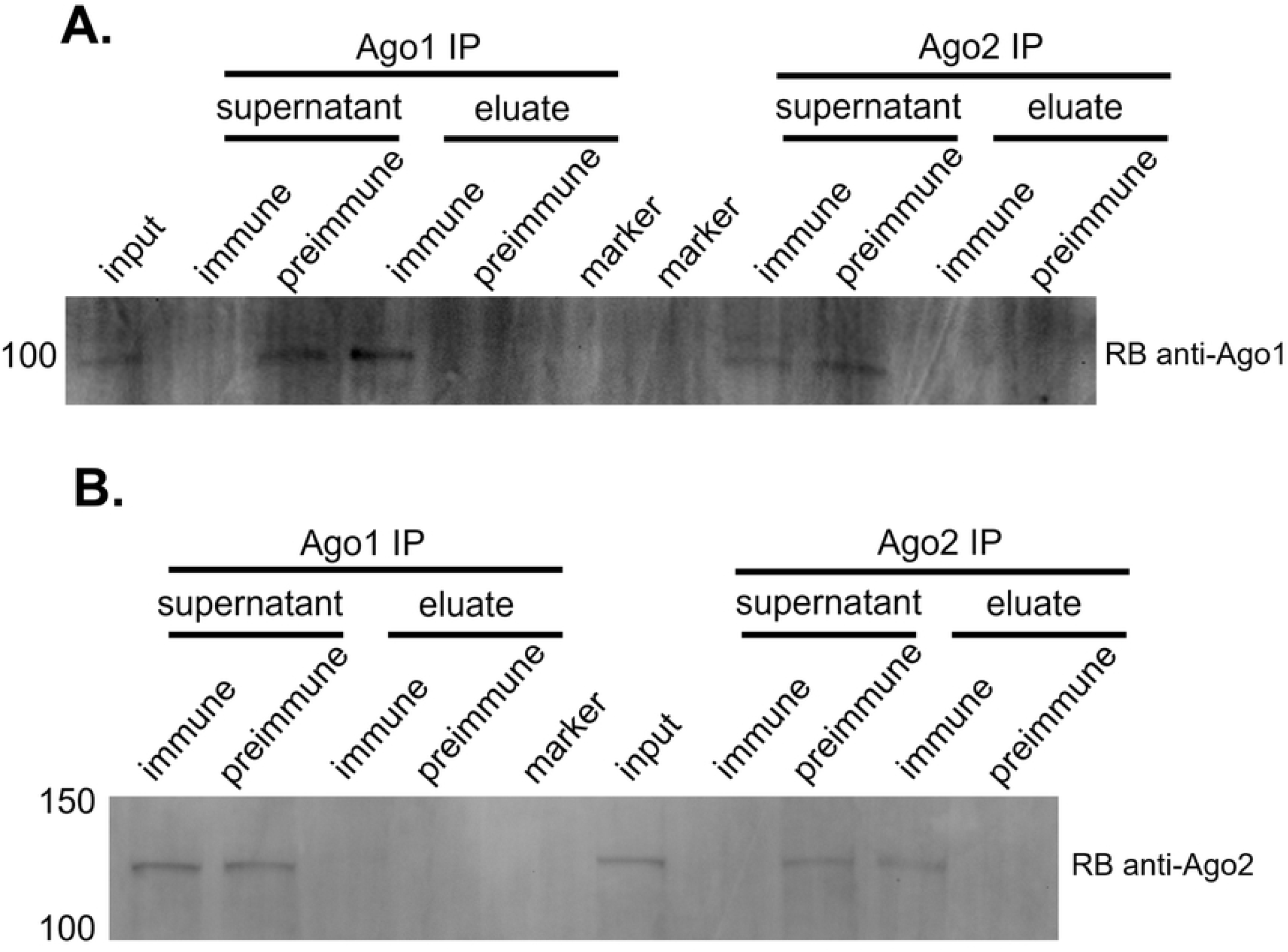
Ago1 and Ago2 antisera effectively and specifically immunoprecipitate Ago1 and Ago2. (A) Guinea pig anti-Ago1 antisera, Guinea pig anti-Ago2 antisera, or preimmune antisera were used in an immunoprecipitation (IP) using lysate from *Drosophila melanogaster* Schneider 2 cells as input. Western blot of input, supernatant, and eluate was probed with a commercial rabbit (RB) anti-Ago1 antibody. Anti-Ago1 antisera immunoprecipitated Ago1 and was capable of depleting the lysate of Ago1. Preimmune sera and anti-Ago2 antisera did not immunoprecipitate Ago1. (B) A second blot was probed with a commercial rabbit anti-Ago2 antibody. Similarly, anti-Ago2 antisera was able to immunoprecipitate Ago2 and depleted the lysate of Ago2. Preimmune sera and anti-Ago1 antisera did not immunoprecipitate Ago2.

**S2 Fig.**
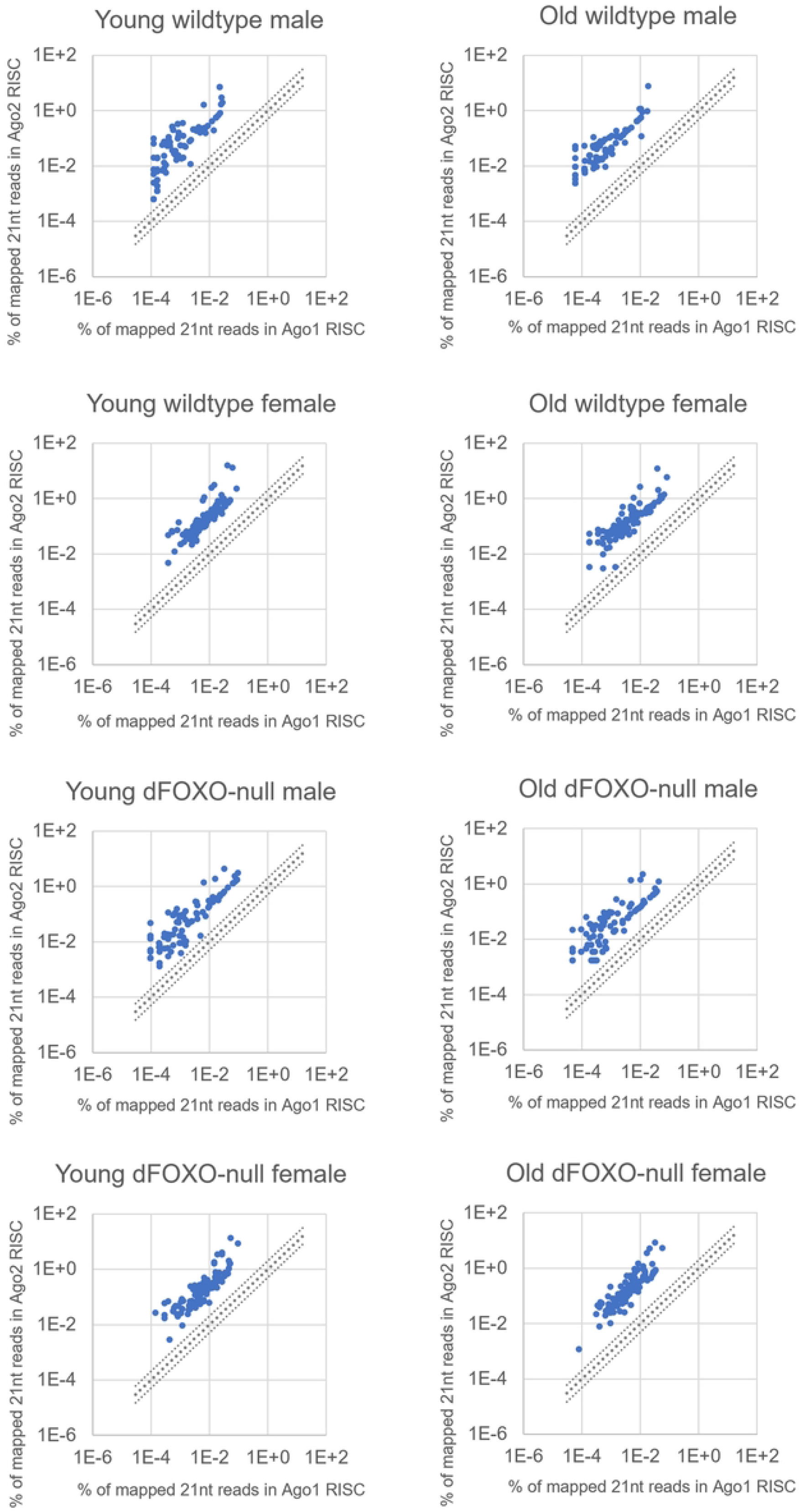
siRNAs against transposons make up larger percentages of Ago2 RISC than Ago1 RISC. Small RNA immunoprecipitated with Ago1 RISC or Ago2 RISC from young (5 day old) or old (35 day old) whole flies were sequenced, size selected to 21 nt *in silico,* and mapped to known *Drosophila* miRNAs and transposons (n = 2). Transposon siRNAs are plotted as percent of mapped reads in Ago1 RISC against percent of mapped reads in Ago2 RISC. All transposon siRNAs that were detected in both RISC make up a greater percentage of Ago2 RISC.

**S1 Table. 20 miRNAs make up the majority of miRNAs detected in total small RNA, Ago1, and Ago2 RISC.** Small RNAs from whole flies, Ago1 RISC immunoprecipitation (IP), or Ago2 RISC IP were sequenced, mapped to known *Drosophila* miRNAs and transposons, and normalized as a percent of total reads mapped (n = 2). miRNA reads were then normalized by total miRNA reads and shown here. (A) The 20 most abundant miRNAs in young wildtype males make up 87 to 94 percent of the total miRNA reads in the male libraries. (B) The 20 most abundant miRNAs in young wildtype females make up 79 to 87 percent of the total miRNA reads in the female libraries.

**S2 Table. 20 transposon siRNAs make up the majority of transposon siRNAs detected in total small RNA and Ago2 RISC.** Small RNAs from whole flies, Ago1 RISC immunoprecipitation (IP), or Ago2 RISC IP were sequenced, size selected in silico, mapped to known *Drosophila* miRNAs and transposons, and normalized as a percent of total reads mapped (n = 2). Transposon siRNA reads were then normalized by total transposon siRNA reads and shown here. (A) The 20 most abundant transposon siRNAs in young wildtype males make up 57 to 88 percent of the total transposon siRNA reads in male total small RNA and Ago2 RISC. (B) The 20 most abundant transposon siRNAs in young wildtype females make up 55 to 66 percent of the total transposon siRNA reads in the female total small RNA and Ago2 RISC.

## Notes

### Competing Interest Statement

The authors have declared no competing interest.

